# Discovery of oxyacanthine dihydrochloride monohydrate polymorphs from obfuscated samples by MicroED

**DOI:** 10.1101/2025.07.03.663078

**Authors:** Jieye Lin, Orel Paz, Johan Unge, Tamir Gonen

## Abstract

It is estimated that more than 50% of marketed pharmaceuticals are derived from natural products. Structural characterization of natural products and their drug formulations is essential for the pharmaceutical industry. Here we report the use of microcrystal electron diffraction (MicroED), to identify two polymorphic crystal structures of oxyacanthine dihydrochloride monohydrate from obfuscated samples that were mislabeled as “berbamine dihydrochloride”. The two polymorphs display primary conformational differences in one of the tetrahydroisoquinoline rings: one polymorph exhibits an intermediate conformation between half-chair and half-boat, while the other adopts a distinct half-boat conformation. Analysis of their structures, energies and crystal packing diagrams indicates a thermodynamic preference for a transformation into the latter. This study highlights the value of integrating MicroED into pharmaceutical pipelines as an efficient tool for structural analysis and quality control.

## 1 Introduction

Natural products exhibit a wide range of biological activities and remain a vital source for drug discovery. It is estimated that more than 50% of marketed pharmaceuticals are derived from natural products.^1,2^ Their diverse biological activities largely originate from structural diversities, including variations in backbone, functional groups, linkages, *etc*.^2^ Elucidating their three-dimensional (3D) atomic structures is fundamental for drug design and development. Traditional techniques such as high-performance liquid chromatography (HPLC), nuclear magnetic resonance (NMR), and high-resolution mass spectrometry (HRMS) are commonly used in combination to determine the chemical structure of natural products.^3^ Single-crystal X-ray diffraction (SC-XRD) remains the gold standard for determining their 3D structures, particularly advantageous for stereochemical assignments. However, it typically requires a well-formed crystal larger than 5 µm,^4^ which is limiting in applications to samples that only form microcrystals, including some natural products.

The cryoEM method microcrystal electron diffraction (MicroED) has demonstrated significant advancements over other approaches to determining pharmaceutical structures.^5,6^ MicroED requires only a billionth of the crystal volume needed for conventional SC-XRD, enabling direct structural determination from powders.^7^ Its rapid data acquisition (*i.e.* 1-2 minutes per datasets) allows for high-throughput analysis within limited instrument time.^8^ MicroED has demonstrated a broad usage in compositional analysis,^8^ impurity detection,^9^ solvomorphism or polymorphism characterization^10-12^ of natural products and their pharmaceutical derivatives.

Berbamine is a bisbenzylisoquinoline alkaloid purified from *Berberis amurensis*, known for biological activities such as antiviral,^13,14^ anti-inflammatory,^15^ *etc*. It is commercially available in both free base and dihydrochloride forms. The free base, which exhibits poor solubility in water, has recently had its crystal structure solved by SC-XRD (CSD entry: PUTZUD).^16^ The Cl^−^ ions in the dihydrochloride form were added to improve water solubility,^25^ yet its structural basis remains unexplored. Oxyacanthine is an isomer of berbamine, displaying similar antiviral activities (*e.g.* SARS-CoV-2).^17^ Their primary structural differences lie in two chiral centers and the position of the phenylhydroxyl group (Figure S1 in Supporting Information). Oxyacanthine is commercially available in free base, dihydrochloride or sulfate forms. The free base structure has been published (CSD entries: HIHKIS, PUTYIQ)^16,18^ but the latter two remain elusive. Berbamine and oxyacanthine are difficult to distinguish from each other due to their similarities in HRMS, HPLC, ^1^H-NMR and ^13^C-NMR and a lack of comprehensive literature demonstration.

Commercial vendors have misidentified these compounds, for example, Cheng *et al.*, (2025) reported the misidentification of oxyacanthine as berbamine by 14 different suppliers.^16^

In this study, two samples labeled as “berbamine dihydrochloride” obtained from different vendors, were subjected to MicroED analysis. Unexpectedly, both were identified as oxyacanthine dihydrochloride monohydrate as the major component. We report here, for the first time, solving two polymorphic structures of oxyacanthine dihydrochloride monohydrate. These polymorphs differ primarily in the tetrahydroisoquinoline (TIQ) ring (*i.e.*, TIQ2 ring in Figure 1): one polymorph exhibits an intermediate TIQ2 ring conformation between half-chair and half-boat, while the other adopts a half-boat TIQ2 ring conformation.^18^ Analysis of their structures, energies and crystal packing diagrams indicates a thermodynamic preference for transformation into the latter. These findings highlight the application of MicroED as a rapid and reliable tool for the structural elucidation, impurity or polymorph detection in natural products, offering valuable guidance for pharmaceutical formulation and quality control.

**Figure 1.**
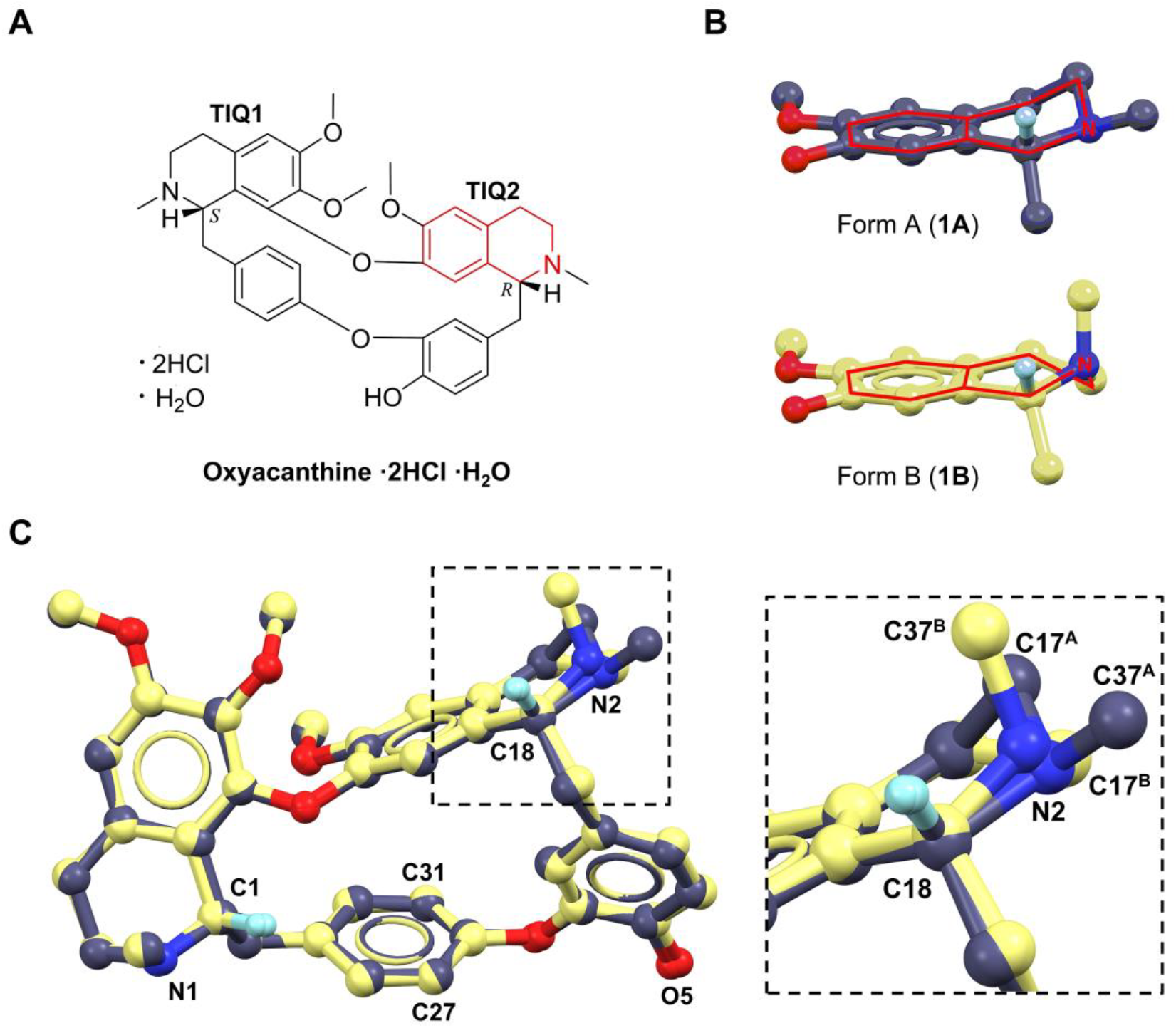
(A) Chemical structure of oxyacanthine dihydrochloride monohydrate; (B) TIQ2 ring distortions in **1A** and **1B**; (C) Overlay of **1A** and **1B** structures showing differences of TIQ2 ring distortions. Cl^−^ ions, water and most of H atoms were omitted for clarification. **1A** is presented in purple, **1B** is presented in yellow.

## 2 Results and discussions

The first “berbamine dihydrochloride” sample was obtained from vendor 1 (V29939; ≥ 98%) and recrystallized from acetone to yield white microcrystals. Crystals were gently scraped into fine powders and transferred onto a TEM grid for MicroED analysis (see details in “Methods”). All microcrystals appeared consistently plate-shaped and exhibited high-resolution diffraction (Figures S2-3 in Supporting Information). A total of 9 crystals were randomly selected, 5 datasets were indexed in the P2_1_ space group with unit cell parameters of **a**=13.60 Å, **b**=9.47Å, **c**=14.66 Å, **β**=115.048°; 4 datasets were indexed in C2 space group with unit cell parameters of **a**=27.94 Å, **b**=9.31 Å, **c**=13.52 Å, **β**=104.531° (Table S1 in Supporting Information). Both unit cell parameters showed close values in **b, c** lengths and **β** angle, while the former has **a** axis length half of the latter. Comparison the picked and predicted spots and spacing measured from their reciprocal spaces validated the correct index (Figures S4-5 in Supporting Information).

These results indicate the co-existence of two polymorphs in an approximate ratio of form A: form B ≈ 5:4. Next, we ordered a second “berbamine dihydrochloride” sample from vendor 2 (HY-N0714A; 96.49%) and conducted MicroED analysis. All microcrystals appeared in the same shape and comparably to the first sample (Figures S2-3 in Supporting Information), therefore a total of 20 crystals were analyzed with no selection preference. 19 datasets were indexed in C2 space group with unit cell parameters of **a**=27.70 Å, **b**=9.29 Å, **c**=13.56 Å, **β**=105.437°, excluding 1 impurity dataset (Table S1 in Supporting Information). We directly solved the MicroED structures by SHELXT/D,^19,20^ revealing that both samples are oxyacanthine dihydrochloride monohydrate, rather than the labeled “berbamine dihydrochloride”. From vendor 1 sample, we solved two polymorphic structures, namely, **1A** (form A) and **1B** (form B). The *ab initio* determined heavier atoms in **1A** and **1B** display a consistent chirality like oxyacanthine. Specifically, the chiral centers at C1 and C18 atoms anticipated to be in *R*- and *S*-configurations in berbamine respectively, are inverted (Figure 1C; Figures S1, S6 in Supporting Information). Additionally, the unique phenylhydroxyl group (O5 atom) is connected to C23 atom rather than C27/C31 atom (Figure 1C; Figure S1, S9 in Supporting Information). The C23−O5 bond lengths were measured as 1.327 Å in **1A** and 1.346 Å in **1B**, closely matching with the typical C_ar_−OH (1.362 ± 0.030 Å) bond length determined by X-ray.^21^ The structures of **1A** and **1B** overlapped well with literature-reported oxyacanthine (free base), with only 0.304 Å and 0.469 Å RMSDs, respectively.^16^ From vendor 2 sample, we only solved **1B** structure which is identical to prior tests on vendor 1 sample.

Statistical analysis of the indexed unit cell parameters supports that **1A** and **1B** are major components in both samples, rather than trace impurities, which is contrary to the high purities (*i.e.* 98% and 96.49%) listed by vendors 1 and 2 (Table S1 in Supporting Information). As reported in literature, oxyacanthine was likely misidentified as berbamine by multiple vendors.^16^ We contacted vendor 1, however they refused to admit the mislabeling according to NMR and MS spectra. Vendor 2 retested the sample by HPLC and NMR, suggesting the sample is pure but it may not correspond to berbamine. Resynthesis of this compound gave similar results. Vendor 2 concluded the misidentification of oxyacanthine as berbamine in that sample.

The crystal structures of **1A** and **1B** were further refined by SHELXL using the electron scattering factors (see details in “Methods”).^22^ H atoms from hydroxyl and amine groups were freely refined from Fo-Fc map, while the remaining H atoms were placed at their geometrically calculated positions. The Cl^−^ ions and water were assigned from analyzing the electron potential maps and the R1 factor in the refinement process. In **1A**, Cl2 ion was placed approximately at position ∼3.18 Å and 2.97 Å away from N2 and O5 atoms, respectively; the minimum Cl^−^ … Cl^−^ distance must be larger than 3.6 Å (calculated from the Cl^−^ radius),^23^ thus a water (OW) was assigned at position ∼2.77 Å near N1 atom, which bridges it with Cl1 and Cl2 ions; in **1B**, two Cl^−^ ions directly bound to amines. The water (OW) bridges two hydrogen bonds between O5 atoms. All the atoms of **1A** and **1B** exhibited a perfect fit to their electrostatic potential maps.

The relatively low R1 values further support the high quality of structural refinements (Figure S7 and Table S2 in Supporting Information).

Superposition of the crystal structures of **1A** and **1B** revealed an excellent consistency (RMSD: 0.35 Å), except for the differences in TIQ2 rings. TIQ1 ring adopts a half-boat conformation for **1A** and **1B**; TIQ2 ring displays an intermediate conformation between a half-boat and a half-chair for **1A,** but in a half-boat conformation for **1B** (Figure 1).^18^ For instance, the N2−C37 bond is equational in **1A** but changes to axial in **1B**; the ring atom C17^A^ is 0.76 Å above the phenyl plane, while the ring atom C17^B^ is 0.30 Å below the plane (Figures 1B-C). The major differences exhibited in C18−N2, C17−N2 and C16−C17 bond rotations of **1A** and **1B** (Figures 2B-C; Figure S8 in Supporting Information). Specifically, C14−C18−N2−C17 rotates from 25.20° to -48.78°, C14−C18−N2−C37 rotates from 159.20° to 78.83°, C16−C17−N2−C18 rotates from -55.27° to 60.66°, C13−C16−C17−N2 rotates from 57.27° to -40.22°. These concerted rotations result in a flip of the N2–C37 bond from an equatorial to an axial orientation, accompanied by a compensatory ring distortion around C17−N2 and C16−C17 bonds.

**Figure 2.**
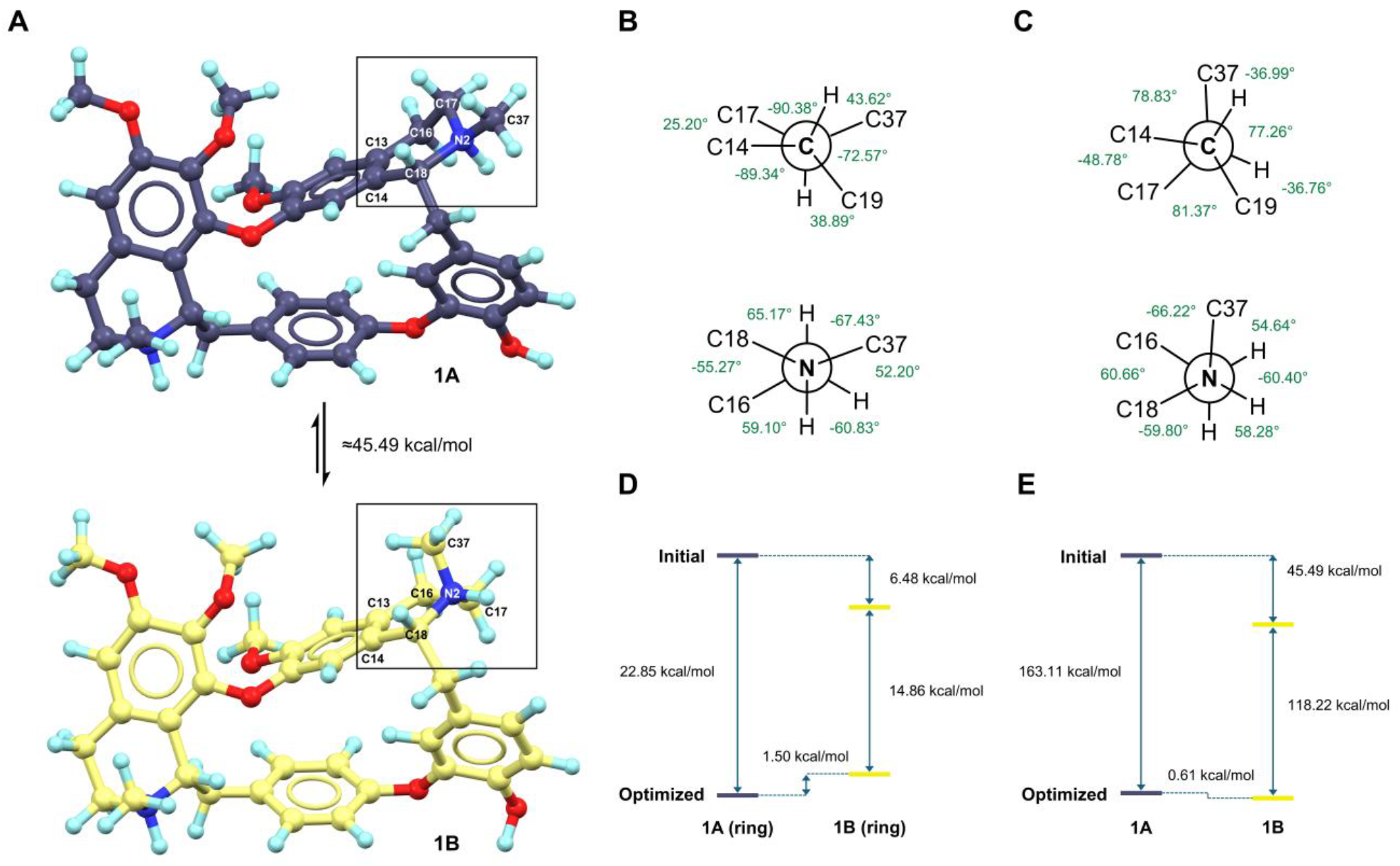
Structural and energy differences between **1A** and **1B**. (A) Crystal structures of **1A** and **1B**; (B) Newman projection of TIQ2 ring in **1A**, viewed along C18−N2 and N2−C17 bonds; (C) Newman projection of TIQ2 ring in **1B**, viewed along C18−N2 and N2−C17 bonds; (D) Single-point energy calculations of initial and geometric-optimized TIQ2 rings; (E) Single-point energy calculations of initial and geometric-optimized **1A** and **1B**. DFT calculations were conducted using *in silico* models A-D (see Figure S8 in Supporting Information).

Neither of the TIQ2 rings in **1A** and **1B** adopts a geometric-idealized conformation. Viewing along the C18−N2 bond (Figures 2B-C), C14 and C17 atoms are near an eclipsed conformation in **1A** (C14−C18−N2−C17 is 15.20°) but are close to a staggered conformation in **1B** (C14−C18−N2−C17 is -48.78°). Viewing along the N2−C17 bond (Figures 2B-C), both C37 and C16 atoms are near staggered conformations in **1A** and **1B**. The former is closer to *anti-* interactions (C16−C17−N2−C37 is 172.13°), while the latter is closer to *gauche*-interactions (C16−C17−N2−C37 is -66.22°).

Single-point DFT energy calculations were performed to evaluate the relative thermodynamic stability of **1A** and **1B**, using models built from their structural coordinates (See details in “Method”; Figure S8 in Supporting Information). During geometric optimization in solution, their TIQ2 rings didn’t flip but fell into corresponding local energy minima, with only 0.61 kcal/mol energy differences between **1A** and **1B** (Figures 2D-E). However, when considering their whole structures in solid-state, the crystal structure of **1B** is 45.49 kcal/mol energy lower than **1A**, where 6.48 kcal/mol is resulted from the TIQ2 ring distortion (Figures 2D-E). These validated the literature-reported ^1^H-NMR observation of doubling peaks (∼7:3),^16^ since both the half-chair and half-boat ring conformations with approximate energies may co-exist in solution. It is plausible that both **1A** and **1B** initially crystalized in ∼1:1 ratio, subsequently **1A** undergoes crystal transformation into a thermodynamically preferred conformation of **1B**.

The point of a crystal transformation is supported by the different crystal packing in **1A** and **1B. 1A** was packed in pairs using a “head-to-head” fashion (Figure 3A; Table S3 in Supporting Information). Two interactive hydrogen bonds N2−H…Cl2 and O5**−**H**…**Cl2 with distances of 3.18 Å and 2.97 Å extend the crystal packing along ***b***-and ***c***-axis. Another Cl1 ion is not directly bound to N1 atom, which is connected by a water (OW) bridge, including three hydrogen bonds, *i.e.*, N1−H…OW (2.77 Å, along ***b***-axis), OW−H…Cl1 (2.93 Å, along ***a***-axis) and OW−H…Cl2 (3.20 Å, along ***a***-axis). The strength of N−H…O is weaker than N−H…Cl^−^ interaction, resulting **1A** being in a higher energy state more susceptible to ion rearrangements. Moreover, the spaces of Cl^−^ ions and water occupied in **1A** are relatively large and continuous (Figure S9A in Supporting Information), which likely to facilitate the rearrangement of Cl^−^ ion and molecule. **1B** was packed in repetitive layer extended by N1**−**H**…**Cl1 (3.07 Å) hydrogen bonding along the ***b***-axis (Figure 3B; Table S3 in Supporting Information). These layers are further connected by interactive hydrogen bonds like N2−H…Cl2 (3.07 Å along ***c***-axis). Additionally, the water (OW) forms two hydrogen bonds with O5 atom (2.67 Å), involving in a repetitive hydrogen bonding chain from OW**−**H**…**O5 to O5**−**H**…**O6 along the ***a***-axis. Compared to **1A**, the spaces where the Cl^−^ ions and water are positioned in **1B** are more separated from each other and more compressed, contributing to a greater structural stability.

**Figure 3.**
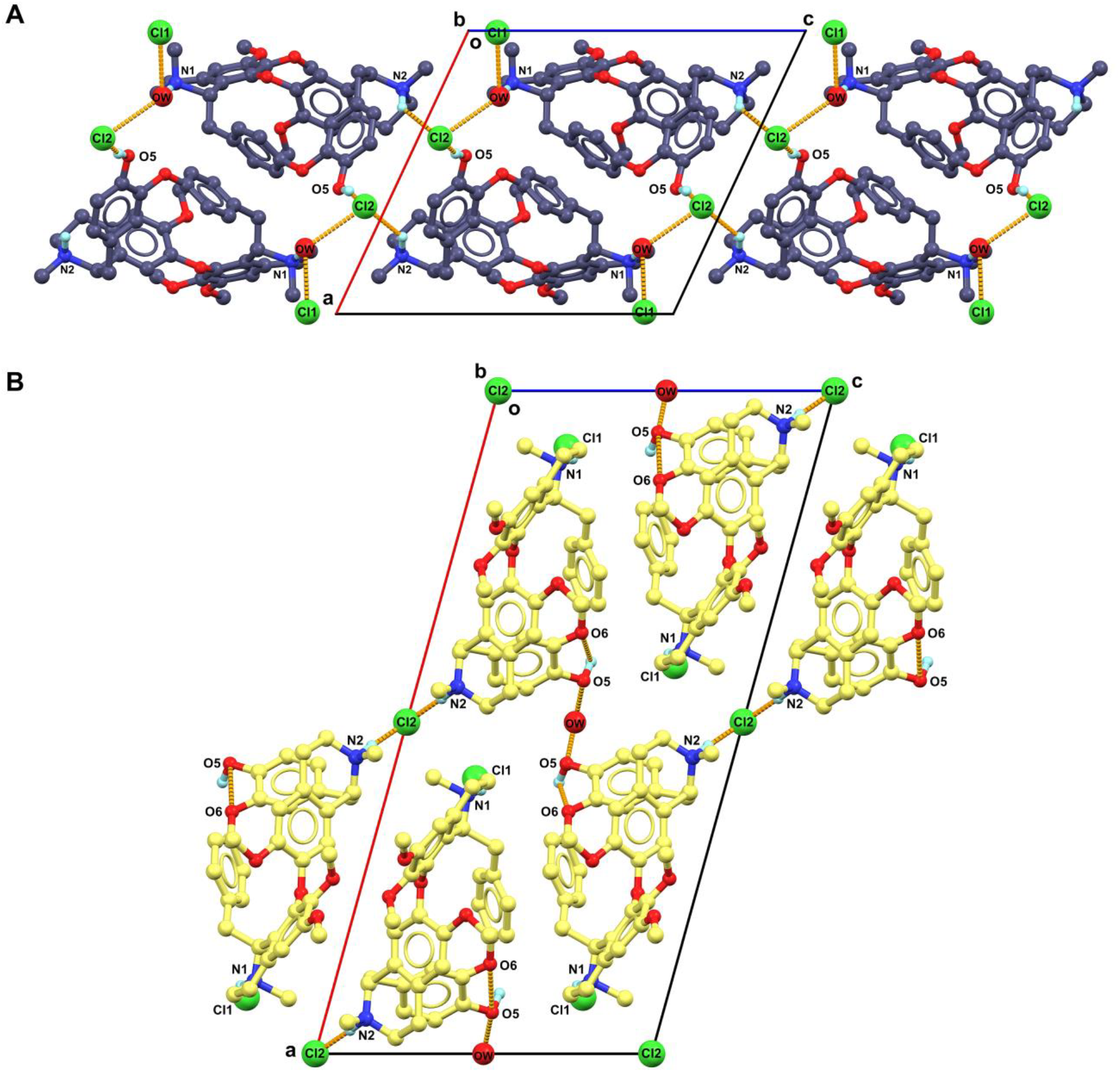
Comparison of different packing diagrams of (A) **1A** and (B) **1B.**Most of H atoms were omitted for clarification.

Oxyacanthine has previously been reported as a potent anti-SARS-CoV-2 inhibitor (IC_50_ = 14.50 µM).^17^ Its binding mechanism with SARS-CoV-2 main protease (abbreviated as “MPro”) has been calculated.^24^ To explore how the polymorphic structures **1A** and **1B** may influence protein-ligand interactions, we docked them into SARS-CoV-2 MPro (PDB entry: 6W63) using the CB-Dock2 web tool. The top-ranked *in silico* models of SARS-CoV-2 MPro-**1A** and SARS-CoV-2 MPro-**1B** closely resemble the literature (Figure 4).^24^ O5 atom in both models serves as a major anchoring site, bridging up to 3 hydrogen bonds with Arg188, Thr199 and Gln192 residues; N1 and O1 atoms can potentially form hydrogen bonds with His41 and Gly143 residues (Figures 4B, 4D). Their major differences lie in the N2 atom. In **1A**, the lone pair (lp) electrons of N2 atom participate in lp…π interaction towards the adjacent phenyl ring, with lp…π *pseudo*-distance of ∼3.24 Å and *pseudo*-angle N2…lp…pi of 134.95°; In contrast, the lp in **1B** positions outside with lp…pi *pseudo*-distance of ∼4.35 Å and *pseudo*-angle N2…lp…pi of 81.21° (Figures 4A, 4C). This outward-facing orientation can potentially be the hydrogen bond acceptor to a water or Glu166 residue (Figure 4D),^18^ thereby enhancing the binding affinity.

**Figure 4.**
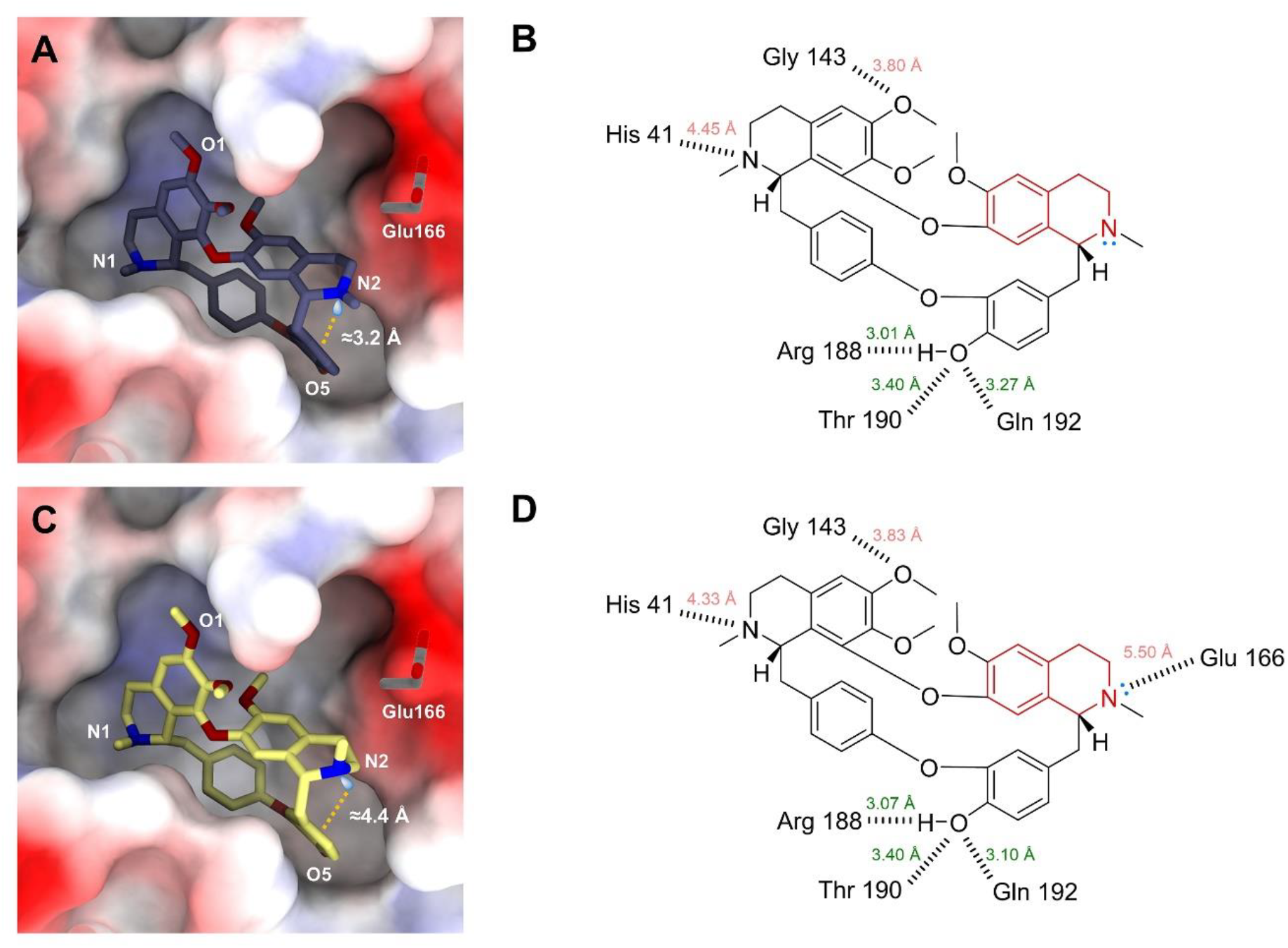
Protein-ligand binding analysis between SARS-CoV-2 MPro (PDB entry: 6W63) and **1A** and **1B.** (A, C) Protein-ligand binding diagram for SARS-CoV-2 MPro-**1A** and SARS-CoV-2 MPro-**1B.** The pseudo-distance between the N2 lone pair electrons and the centroid of phenyl ring is displayed in orange. (B, D) Potential hydrogen bonding interactions between **1A**/**1B** and SARS-CoV-2 MPro.

## 3 Conclusion

Formulating a natural product candidate for the pharmaceutical market requires substantial efforts. For example, salt formulation is commonly employed to enhance the solubility and bioavailability of hydrophobic compounds.^25^ In quality control, impurity and polymorph analyses are critical and typically rely on multiple techniques such as NMR, SC-XRD, *etc*. Failure to detect or control these factors can lead to significant commercial losses, as exemplified by the recall of Norvir (an HIV protease inhibitor) in 1998 due to unexpected polymorphic contamination.^26^

In this study, we demonstrated an efficient application of MicroED in identifying impurities. In this case, mislabeled “berbamine dihydrochloride” samples as oxyacanthine dihydrochloride monohydrate which was originally misidentified.^16^ The success rate of solving structures of pharmaceuticals with MicroED is extremely high compared with SXRD and often the attainable resolution may also be better.^27^ MicroED enables a rapid and direct analysis of powder mixtures, requiring only 1-2 minutes per dataset for data collection and minutes to determine a definitive crystal structure.^7^ These findings highlight the value of incorporating MicroED into pharmaceutical formulation pipelines for structural analysis and quality control.

From a structural perspective, two polymorphs **1A** and **1B** were firstly resolved from the mixture, differing primarily in their TIQ2 ring conformations. This discovery refreshed our understanding of oxyacanthine, as it represents the first-time observation of a half-boat conformation in TIQ2 ring of **1B** in oxyacanthine’s dihydrochloride form, distinct from both **1A** and previously known free base structures.^16,18^ It is assumed that such polymorphism is caused by crystal transformation where a relocation of the Cl^−^ ion and molecules occur. Energy calculations and crystal packing analyses support it showing a thermodynamic preference for **1B** over **1A**. Notably, the lone pair electron of N2 atom in **1B** is positioned outside, potentially can form additional hydrogen bond to enhance the protein-ligand binding,^18^ whereas it is absent in both **1A** and free base forms. These findings indicate that **1B** might be more suitable for pharmaceutical production, although the continued monitoring for the presence of **1A** remains essential during production.

## 4 Methods

### Sample preparation

Two samples labeled as “berbamine dihydrochloride” were commercially purchased from vendor 1 (V29939; ≥ 98%) and vendor 2 (HY-N0714A; 96.49%). Around 5 mg powders each were recrystallized from acetone with free evaporation in a 10 mL scintillation vial. A spatula was used to scrape crystals from vial resulting in seemingly amorphous powdery sample. Powders were mixed with a 400-mesh glider grid coated with continuous carbon film (3.05 mm O.D., Ted Pella Inc.) which was pretreated with 15 mA negative glow-discharge plasma for 30 s using PELCO easiGlow (Ted Pella Inc.). The grid was clipped outside using c-ring and autogrid ring at room temperature and then loaded into a Thermo Fisher Talos Arctica Cryo-TEM (200 kV, ∼0.0251 Å) equipped with a CMOS CetaD camera (4096 × 4096 pixels).

### MicroED data collection

Manual data collection was conducted in EPU-D software (Thermo Fisher).^7^ The microcrystals were screened under imaging mode (SA 3400×), where only the thin crystals with a light visually contrast to the carbon film were selected. Their eucentric heights were calibrated to maintain the crystals inside the beam during the continuous rotation. MicroED data was collected in parallel illumination in diffraction mode (659 mm), spot size 11 (µP), 70 µm C2 aperture and 100 µm selected area (SA) aperture resulting in ∼0.01 e^-1^/Å^2^/s dose rate.^28^ The stage was constantly rotated at 1 °/s over an angular wedge of 120° from -60° to +60°, with 1 s exposure time per frame, resulting a total dose of ∼1.20 e^-^/Å^2^ for each dataset.

### MicroED data processing

MicroED data was saved in mrc format and converted to smv format using the mrc2smv software (https://cryoem.ucla.edu/microed).^29^ Images were imported to XDS for spots picking, index and integration.^30,31^ The best datasets were scaled by XSCALE,^31^ converted to SHELX hkl format by XDSCONV,^31^ and *ab initio* solved by SHELXT/D.^19,20^ Their structures were refined by SHELXL^22^ using Shelxle^32^ as a graphical interference to yield the final MicroED structures (Figure 1C, Table S2 in Supporting Information). The H atoms of hydroxyl and amine groups were refined from corresponding Fo-Fc map. All the other H atoms were refined by geometrically calculated positions, with C−H =1.120 (methine), 1.110 (methylene),1.080 (methyl) or 1.100 Å (aryl).^22^

### DFT calculations

DFT calculations were conducted by ωB97X/6-311G(d,p)^33,34^ functional/basis set implemented in ORCA 5.0 software.^35^ DFT models A-D were built based on the coordinates extracted from **1A** and **1B** (Figure S8 in Supporting Information). Solvent effects of water were treated by the conductor-like polarizable continuum model (CPCM)^36^ and the solvation model based on density (SMD).^37^ The resulting single-point energies were converted to kcal/mol and compared to evaluate the relative energy differences (Figures 2D-E).

### Molecular docking

The SARS-CoV-2 MPro structure was retrieved from PDB entry 6W63. The natively bound ligand X77 and water were removed using Pymol.^38^ Ligand structures were built by **1A** and **1B** without Cl^−^ ions, water and the charges in amines. The molecular docking was automatically performed using CB-Dock2 (Cavity-detection guided Blind Docking) web tool.^39^ The docking center was positioned at (-23, 15, -30), consistent with the position of X77 in the original PDB structure,^24^ and the grid box was set as (23, 23, 30). The docking model with the highest binding affinity was analyzed. The protein-ligand binding was analyzed using PLIP (Protein-Ligand Interaction Profiler) web tool and visualized in ChimeraX software (Figure 4).^40,41^

## Acknowledgements

This study was supported by the National Institutes of Health P41GM136508. Portions of this research or manuscript completion were developed with funding from the Department of Defense MCDC-2202-002. Effort sponsored by the U.S. Government under Other Transaction number W15QKN-16-9-1002 between the MCDC, and the Government. The US Government is authorized to reproduce and distribute reprints for Governmental purposes, notwithstanding any copyright notation thereon. The views and conclusions contained herein are those of the authors and should not be interpreted as necessarily representing the official policies or endorsements, either expressed or implied, of the U.S. Government. The PAH shall flow down these requirements to its sub awardees, at all tiers. The Gonen laboratory is supported by funds from the Howard Hughes Medical Institute.

