## Supplemental files for "Discovery of oxyacanthine dihydrochloride monohydrate polymorphs from obfuscated samples by MicroED"

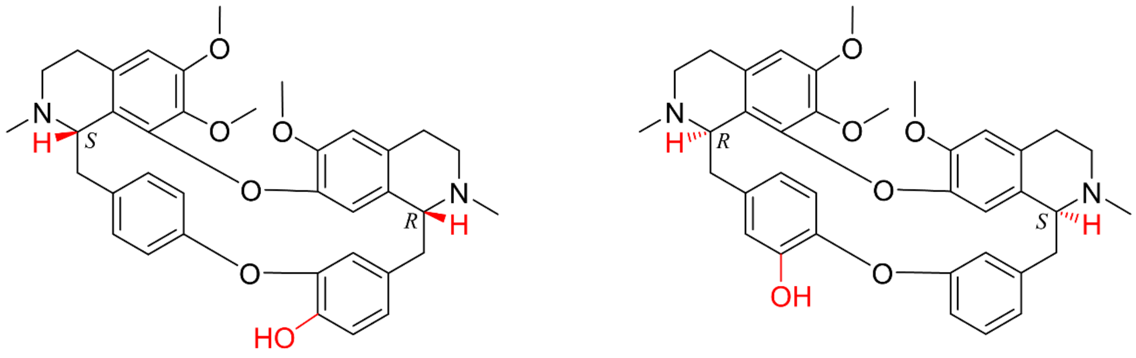

**Oxyacanthine** **Berbamine**

**Figure S1** Chemical structures of oxyacanthine and berbamine showing differences in chiral centers and phenylhydroxyl group.

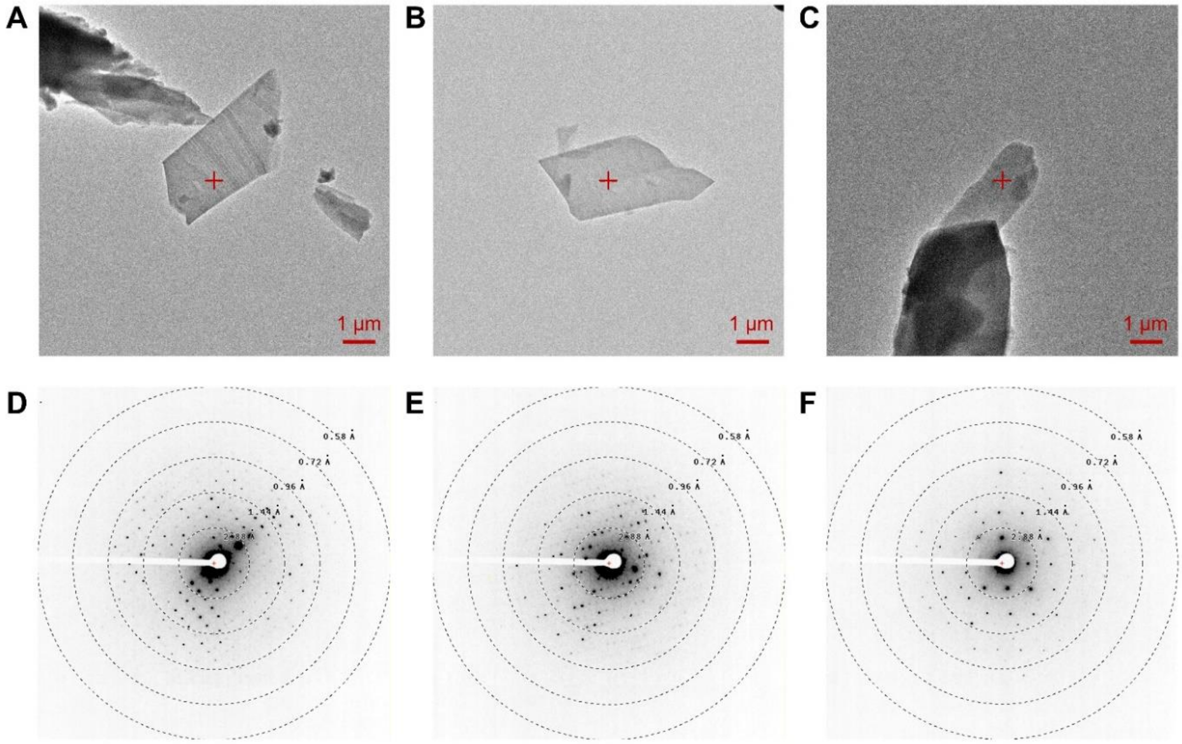


**Figure S2** Crystal appearance and diffraction pattern under the TEM. (A-B) Images of **1A** and **1B** (vendor 1) under the imaging mode (SA 5300×). (C) Image of **1B** (vendor 2) under the imaging mode (SA 5300×). (D-E) Diffraction pattern of **1A** and **1B** (vendor 1) under diffraction mode (659 mm). (F) Diffraction pattern of **1B** (vendor 2) under diffraction mode (659 mm).

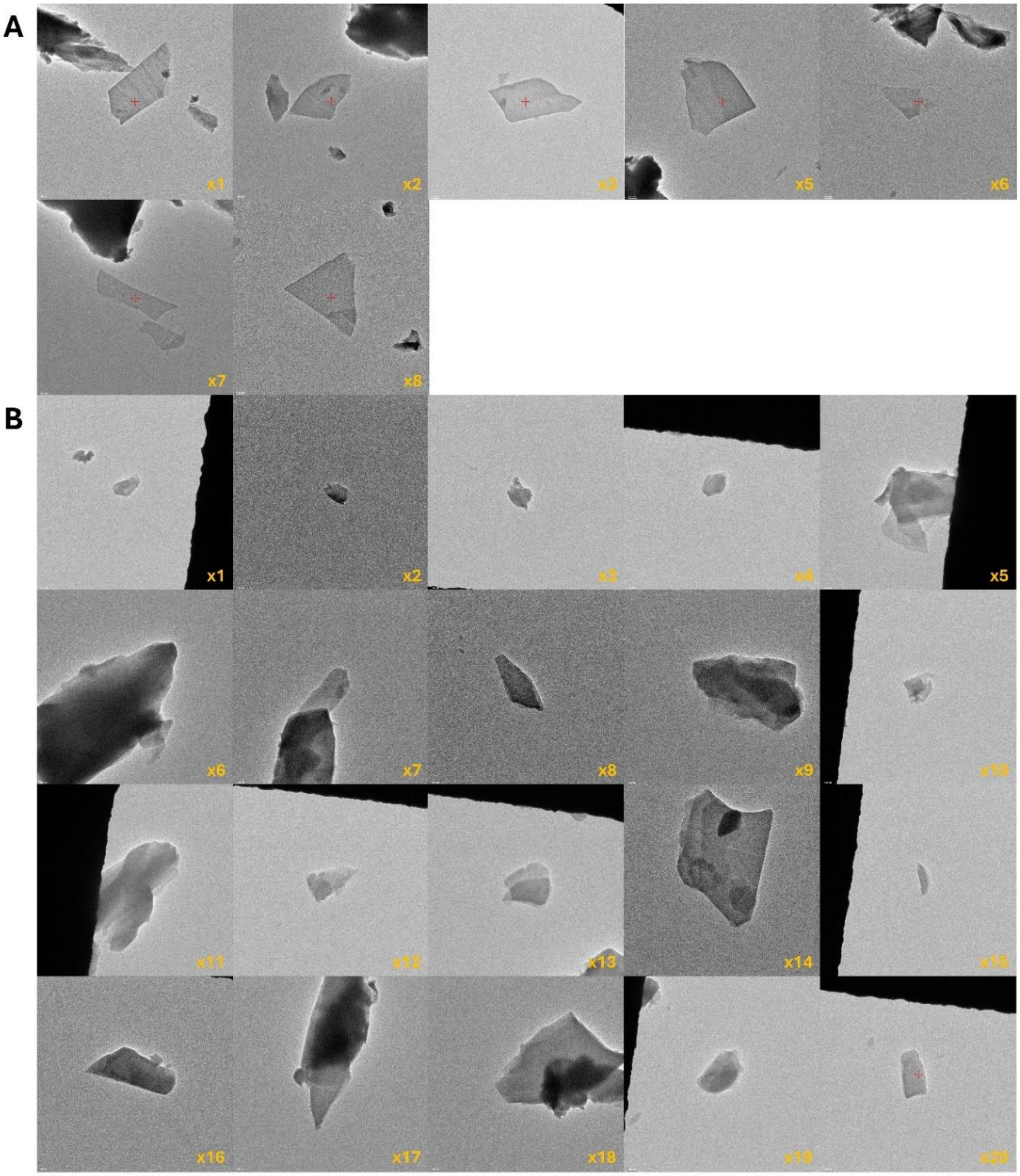


**Figure S3** Crystal appearances of "berbamine dihydrochloride" samples obtained from vendors 1 (A) and 2 (B).

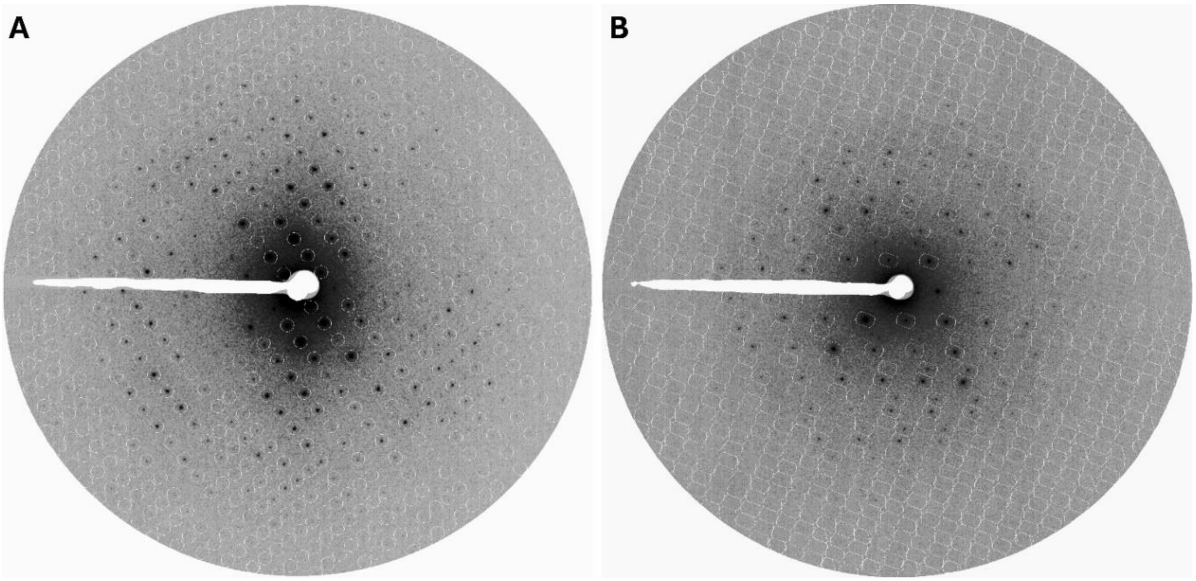


**Figure S4** Comparison of the picked and predicted spots suggesting satisfactory index and integration for (A) **1A** and (B) **1B**.

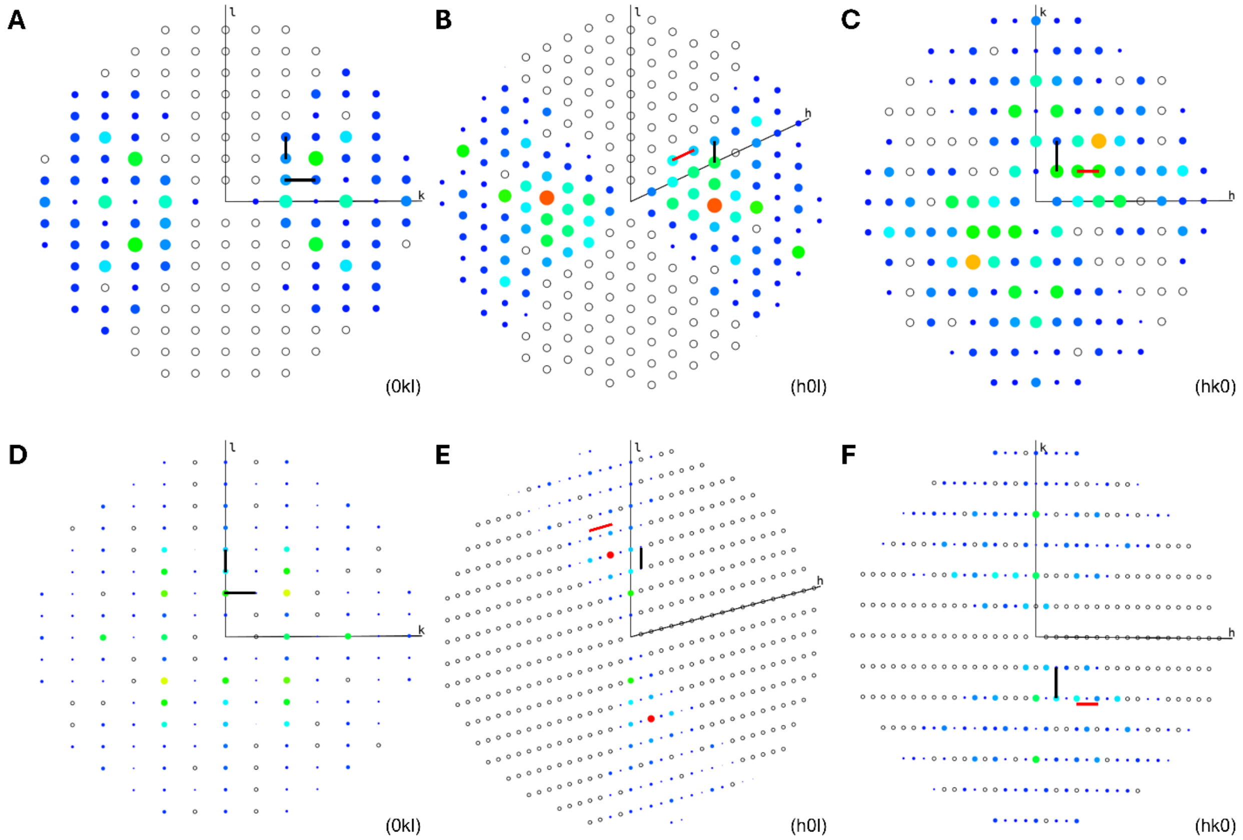


**Figure S5** Reciprocal space of (A-C) **1A** and (D-F) **1B**. Reflections were integrated with P1 space group and the resolution was cut at 1.5 Å.

**A**

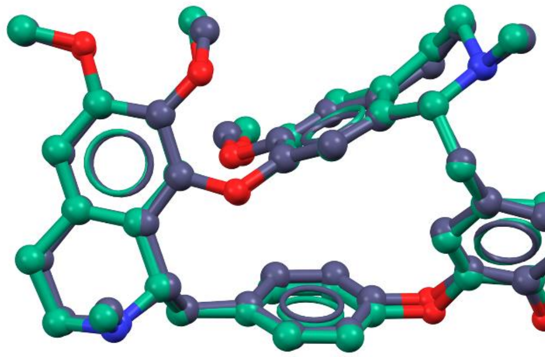

**RMSD:** 0.304 Å

**B**

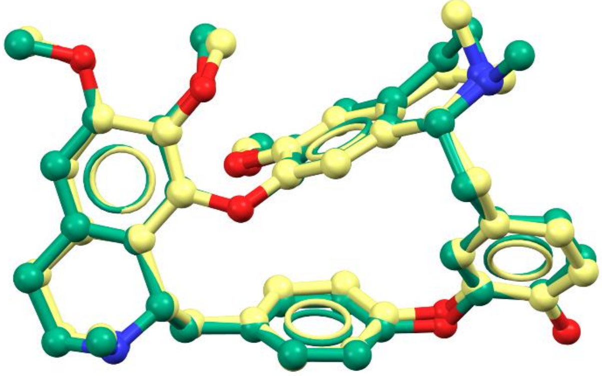

**RMSD:** 0.469 Å

**Figure S6** Superposition of (A) **1A** and (B) **1B** to the literature-reported oxyacanthine (free base) structure.[1] $Cl^-$ ions, water and hydrogen atoms were omitted for clarification. **1A** is colored in purple, **1B** is colored in yellow, the literature-reported oxyacanthine (free base) structure is colored in green.

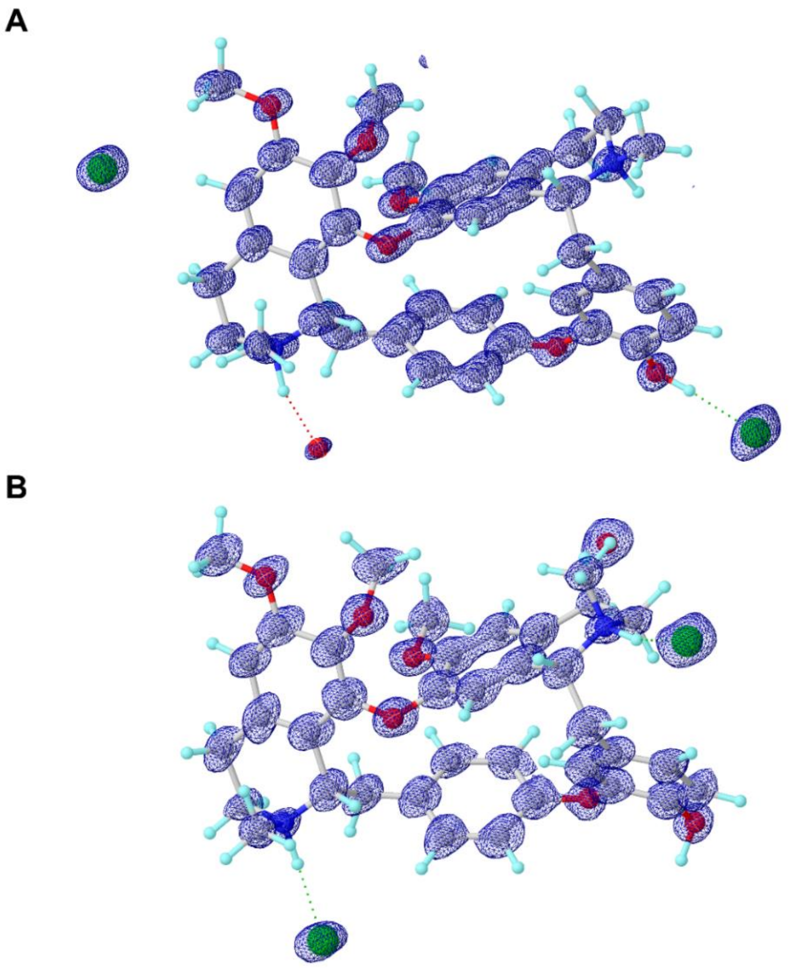


**Figure S7** 2Fo-Fc maps of **1A** and **1B** (contour level: 3σ)

**Model A**

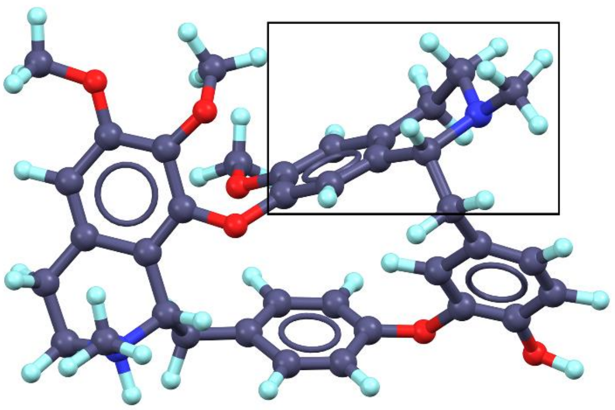

**Model B**

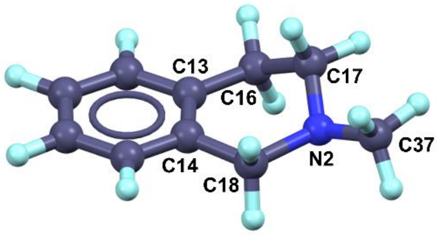


**Torsion angles (°)**

| | | | |
|---|---|---|---|
| C14–C13–C16–C17 | -28.95 | N2–C18–C14–C13 | 6.29 |
| C13–C16–C17–N2 | 57.27 | C18–C14–C13–C16 | -2.76 |
| C16–C17–N2–C18 | -55.27 | C14–C18–N2–C37 | 159.20 |
| C17–N2–C18–C14 | 25.20 | C16–C17–N2–C37 | 172.13 |

**Model C**

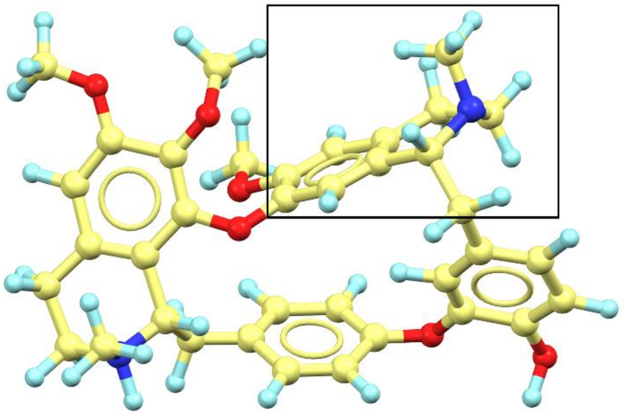

**Model D**

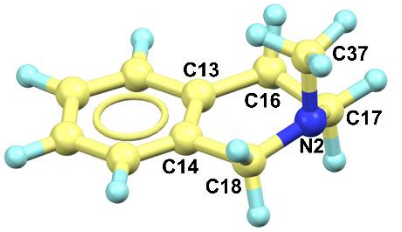


**Torsion angles (°)**

| | | | |
|---|---|---|---|
| C14–C13–C16–C17 | 13.74 | N2–C18–C14–C13 | 16.71 |
| C13–C16–C17–N2 | -40.22 | C18–C14–C13–C16 | -1.58 |
| C16–C17–N2–C18 | 60.66 | C14–C18–N2–C37 | 78.83 |
| C17–N2–C18–C14 | -48.78 | C16–C17–N2–C37 | -66.22 |

**Figure S8** *In silica* models used in DFT calculations. Models A-D were built using the coordinates from **1A** and **1B**.

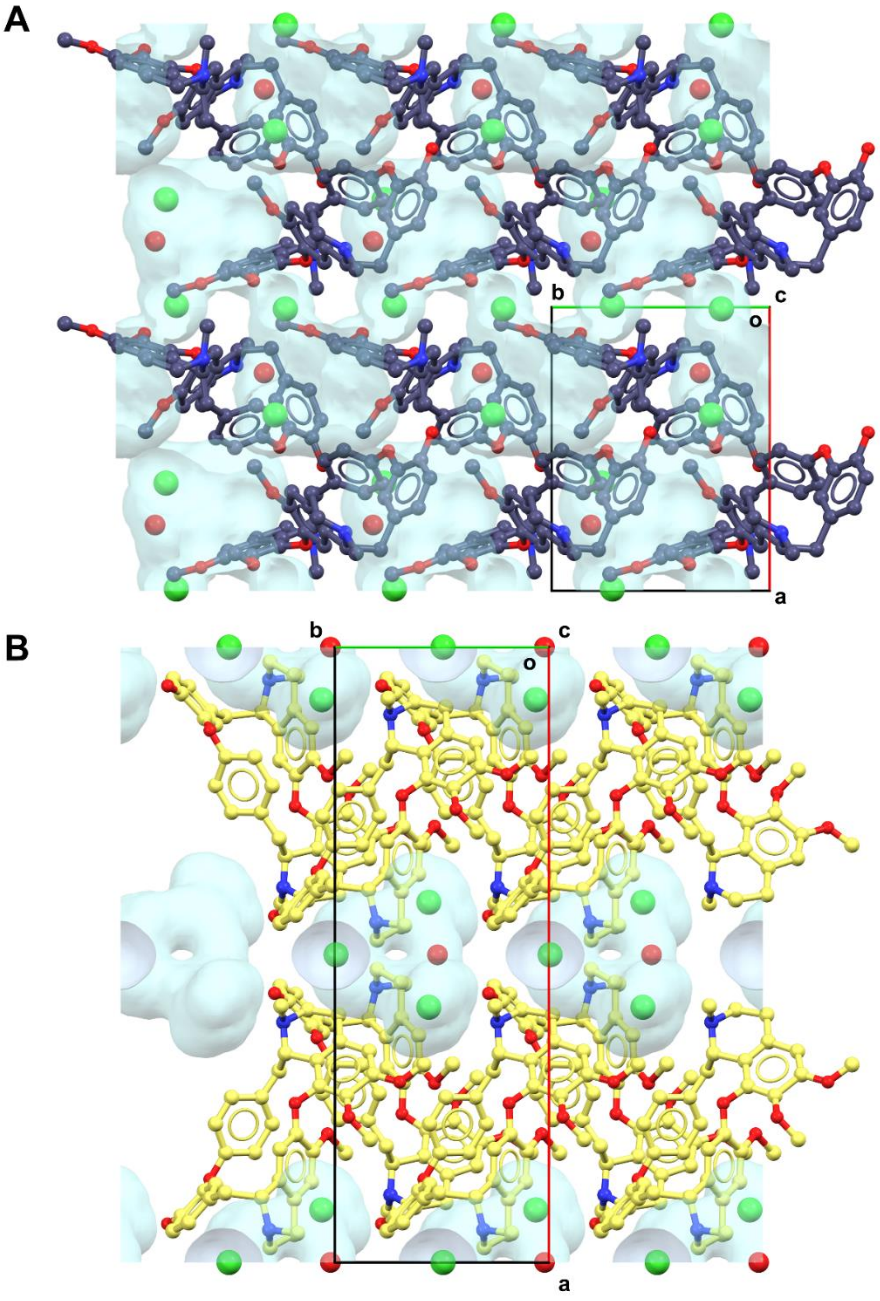


**Figure S9** Contact surface (voids) detected in (A) **1A** and (B) **1B** after removal of $Cl^{-}$ ions and water. Probe radius is 1.2 Å with 0.3 Å approximately grid spacing.

**Table S1** Statistical analysis of the unit cell parameters (Å, °) indexed from vendors 1 and 2.

| Sample | xtal (#) | Space group | **a** | **b** | **c** | **α** | **β** | **γ** |
|---|---|---|---|---|---|---|---|---|
| vendor 1 | 1 | $P2_1$ | 13.60 | 9.47 | 14.65 | 90 | 115.051 | 90 |
| | 2 | $P2_1$ | 13.57 | 9.43 | 14.64 | 90 | 114.642 | 90 |
| | 3 | C2 | 27.94 | 9.31 | 13.52 | 90 | 104.531 | 90 |
| | 4 | C2 | 27.93 | 9.30 | 13.59 | 90 | 105.316 | 90 |
| | 5 | C2 | 27.90 | 9.35 | 13.54 | 90 | 104.787 | 90 |
| | 6 | $P2_1$ | 13.64 | 9.44 | 14.85 | 90 | 115.439 | 90 |
| | 7 | $P2_1$ | 13.61 | 9.49 | 14.57 | 90 | 115.180 | 90 |
| | 8 | $P2_1$ | 13.70 | 9.41 | 15.27 | 90 | 114.043 | 90 |
| | 9 | C2 | 27.62 | 9.37 | 13.62 | 90 | 105.049 | 90 |
| Sample | xtal (#) | Space group | **a** | **b** | **c** | **α** | **β** | **γ** |
| vendor 2 | 1 | C2 | 27.85 | 9.22 | 13.67 | 90 | 105.327 | 90 |
| | 2 | C2 | 27.81 | 9.28 | 13.59 | 90 | 105.630 | 90 |
| | 3 | P1 | 7.06 | 7.42 | 7.56 | 115.134 | 107.137 | 100.337 |
| | 4 | C2 | 27.82 | 9.30 | 13.57 | 90 | 105.664 | 90 |
| | 5 | C2 | 27.75 | 9.30 | 13.54 | 90 | 105.493 | 90 |
| | 6 | C2 | 27.68 | 9.28 | 13.64 | 90 | 105.094 | 90 |
| | 7 | C2 | 27.7 | 9.29 | 13.56 | 90 | 105.437 | 90 |
| | 8 | C2 | 27.63 | 9.26 | 13.60 | 90 | 105.264 | 90 |
| | 9 | C2 | 27.73 | 9.28 | 13.61 | 90 | 105.556 | 90 |
| | 10 | C2 | 28.18 | 9.27 | 13.56 | 90 | 105.744 | 90 |
| | 11 | C2 | 27.61 | 9.29 | 13.69 | 90 | 105.432 | 90 |
| | 12 | C2 | 27.75 | 9.29 | 13.55 | 90 | 105.375 | 90 |
| | 13 | C2 | 27.70 | 9.28 | 13.55 | 90 | 105.084 | 90 |
| | 14 | C2 | 27.49 | 9.39 | 13.56 | 90 | 104.617 | 90 |
| | 15 | C2 | 27.74 | 9.25 | 13.62 | 90 | 105.863 | 90 |
| | 16 | C2 | 27.49 | 9.25 | 13.65 | 90 | 105.400 | 90 |
| | 17 | C2 | 27.68 | 9.26 | 13.62 | 90 | 105.638 | 90 |
| | 18 | C2 | 27.55 | 9.26 | 13.61 | 90 | 104.998 | 90 |
| | 19 | C2 | 27.89 | 9.28 | 13.56 | 90 | 105.24 | 90 |
| | 20 | C2 | 27.92 | 9.24 | 13.59 | 90 | 105.398 | 90 |

**Notes**: The vendor 2 xtal 3 is an impurity because the cell volume is too small for oxyacanthine or berbamine.

**Table S2** MicroED data statistics of **1A** and **1B**.

| | **1A** | **1B** |
|---|---|---|
| Chemical formula | $C_{37}H_{44}Cl_2N_2O_6$ | $C_{37}H_{44}Cl_2N_2O_6$ |
| Molar mass | 697.63 | 697.63 |
| Temperature (K) | 80 | 80 |
| Crystal system | Monoclinic | Monoclinic |
| Space group | $P2_1$ | C2 |
| Unit cell lengths (Å) | | |
| **a** | 13.600 | 27.700 |
| **b** | 9.470 | 9.290 |
| **c** | 14.660 | 13.560 |
| Unit cell angles (°) | | |
| α | 90.00 | 90.00 |
| β | 115.05 | 105.44 |
| γ | 90.00 | 90.00 |
| Cell volume (Å$^3$) | 1710.5 | 3363.5 |
| No. of merged datasets | 1 | 2 |
| No. of observed reflections | 8735 | 11121 |
| No. of unique reflections | 3027 | 2826 |
| $R_{obs}$ (%) | 13.2 | 18.6 |
| $R_{meas}$ (%) | 16.4 | 21.5 |
| I/Sigma | 5.42 | 5.69 |
| $CC_{1/2}$ | 98.2 | 98.8 |
| **Resolution (Å)** | **0.80** | **0.86** |
| **Completeness (%)** | **81.2** | **95.6** |
| **$R_1$ (%)** | **19.28** | **17.19** |
| $wR_2$ (%) | 37.88 | 37.34 |
| GooF | 1.185 | 1.161 |

**Table S3** Hydrogen bonding interactions of **1A** and **1B** (Å, °).

| **1A** | **D–H** | **H···A** | **D···A** | **D–H···A** |
|---|---|---|---|---|
| N1–H···OW | 1.118 | 1.740 | 2.774 | 151.39 |
| N2–H···Cl2 | 1.122 | 2.304 | 3.176 | 132.92 |
| O5–H···Cl2 | 1.000 | 1.994 | 2.972 | 165.40 |
| OW–H···Cl1 | – | – | 2.932 | – |
| OW–H···Cl2 | – | – | 3.201 | – |
| **1B** | **D–H** | **H···A** | **D···A** | **D–H···A** |
| N1–H···Cl1 | 1.124 | 2.074 | 3.065 | 145.28 |
| N2–H···Cl2 | 1.123 | 1.943 | 3.034 | 162.97 |
| O5–H···O6 | 1.001 | 2.728 | 2.671 | 76.16 |
| OW–H···O5 | – | – | 2.801 | – |
